# Multidirectional digital scanned light-sheet microscopy enables uniform fluorescence excitation and contrast-enhanced imaging

**DOI:** 10.1101/270207

**Authors:** Adam K. Glaser, Ye Chen, Chengbo Yin, Linpeng Wei, Lindsey A. Barner, Nicholas P. Reder, Jonathan T.C. Liu

## Abstract

Light-sheet fluorescence microscopy (LSFM) has emerged as a powerful method for rapid and optically efficient 3D microscopy. Initial LSFM designs utilized a static sheet of light, termed selective plane illumination microscopy (SPIM), which exhibited shadowing artifacts and deteriorated contrast due to light scattering. These issues have been addressed, in part, by multidirectional selective plane illumination microscopy (mSPIM), in which rotation of the light sheet is used to mitigate shadowing artifacts, and digital scanned light-sheet microscopy (DSLM), in which confocal line detection is used to reject scattered light. Here we present a simple passive multidirectional digital scanned light-sheet microscopy (mDSLM) architecture that combines the benefits of mSPIM and DSLM. By utilizing an elliptical Gaussian beam with increased angular diversity in the imaging plane, mDSLM provides shadow-free contrast-enhanced imaging of fluorescently labeled samples.

**One Sentence Summary:** Glaser *et al.* describe a light-sheet microscopy architecture that enables passive multidirectional illumination with confocal line detection to enable both uniform fluorescence excitation and contrast-enhanced imaging of fluorescently labeled samples.

## Introduction

Light-sheet fluorescence microscopy (LSFM), whose technological roots may be traced back over a century [1], has recently seen intense development for a wide array of research investigations and potential clinical applications [2–15]. The success of LSFM has stemmed from its ability to achieve extremely high-speed 3D imaging through camera-based detection in a configuration that is more optically efficient and “gentle” than other optical-sectioning approaches in terms of light dose (minimizing photodamage and photobleaching) [11, 16]. The LSFM approach achieves optical sectioning (rejection of out-of-focus light) by exciting fluorescence along a thin 2D illumination “light sheet” within a sample, which is imaged in the orthogonal direction with a high-speed detector array. The flexibility of this “dual-axis” configuration, where the illumination and collection beam paths are decoupled and may be individually optimized, is in contrast to conventional single-axis microscopes in which the illumination and collection beams travel along a common path.

The original LSFM design utilized a static light-sheet architecture and was termed “selective plane illumination microscopy (SPIM)” [9]. While simple and straightforward, this illumination method has a few limitations. First, the lack of angular diversity in the light sheet (i.e. the photons all travel in roughly the same direction), results in shadowing artifacts within the sample due to occlusions [17]. Second, the illumination light sheet is scattered in biological tissues, which generates an unwanted background that reduces image contrast (defined here as signal-to-background ratio, SBR), and consequently, imaging depth. To address the issue of shadowing artifacts, the multidirectional selective plane illumination microscopy (mSPIM) architecture was devised [18], in which the light sheet is rotated in the plane of the sheet to average out the shadowing artifacts over time (assuming that the rotation is faster than the integration time of the detector array). At around the same time that mSPIM was developed, digital scanned light-sheet microscopy (DSLM) with confocal line detection was also developed to enhance image contrast with LSFM [5, 19, 20]. With DSLM, a Gaussian pencil beam is laterally scanned to create a 2D light sheet over time. The scanned pencil beam can be synchronized to the rolling shutter of a sCMOS detector array, which serves as a digital confocal slit to reject out-of-focus scattered light and thereby improve image contrast and/or depth. Unfortunately, mSPIM and confocal DSLM are incompatible since rotating a pencil beam would cause much of the beam to rotate out of the confocal slit. In addition, an exceedingly high rotational rate (two orders of magnitude faster than mSPIM) would be required to match the integration time of a confocal rolling shutter (see Supplementary Materials for details).

While the issues of shadowing and reduced image contrast in the original SPIM design have been independently addressed, in part, by mSPIM and DSLM, a solution that simultaneously addresses both issues has not been reported. Here, we present an approach, termed multidirectional digitally scanned light-sheet microscopy (mDSLM), which utilizes an elliptical Gaussian pencil beam that provides a similar degree of “angular diversity” compared with mSPIM, for mitigation of shadowing artifacts, but does not require rotation of the beam. Since mDLSM is a passive approach, it allows for confocal line detection to achieve improved contrast in comparison to SPIM/mSPIM. Finally, unlike computational approaches [21, 22], mDSLM intrinsically enhances image quality and does not require downstream processing of notoriously large LSFM datasets [23].

Given the growing interest in LSFM for both fundamental and clinical research, the improved image quality provided by the mDLSM approach should enable improved biological investigations as well as higher-fidelity pathology for accurate prognostication and treatment stratification [10, 24, 25].

## Results

### Theory

Although an array of illumination beam types have been explored for LSFM [26–29], including propagation-invariant Bessel and Airy beams with shadow-mitigation properties, the majority of LSFM systems utilize Gaussian beams, for which the intensity is described by a solution to the paraxial Helmholtz equation (see Discussion section for a summary of non-Gaussian beam types). For a Gaussian beam propagating along the *z*-axis, the spatial intensity distribution in Cartesian coordinates (rather than the more-commonly used polar coordinate system), *I*(*x*,*y*,*z*), is given as:

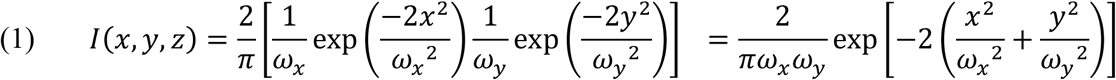

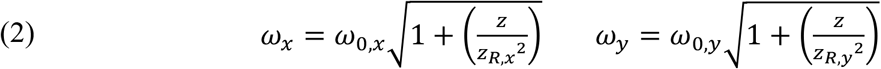

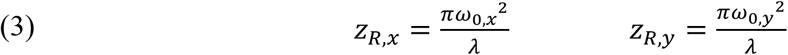

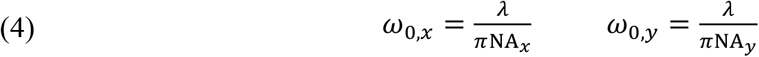

As expressed in Eq. (2), and *ω*_*x*_ and *ω*_*y*_ are the beam radii in the *x* and *y* dimensions respectively (defined at the 1/*e*^2^ intensity points). The Rayleigh ranges in the *x* and *y* dimensions, *z*_*R,x*_ and *z*_*R,y*_, are given by in Eq. (3) and are defined as the axial extent from the beam focus to the point at which the beam radii has expanded to 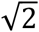 larger than the beam waists, *ω*_0,*x*_ and *ω*_0,*y*_, as expressed in Eq. (4). NA_*x*_ and NA_*y*_ are the numerical aperture (NA) of the beam in the *x* and *y* dimensions, respectively.

With SPIM (Fig. 1a), a cylindrical lens is used to illuminate the back focal plane (BFP) of an infinity-corrected objective (i.e. the Fourier plane) with a line focus, such that a two-dimensional (2D) light sheet is generated within the specimen at the front focal plane (FFP) of the objective. With respect to the coordinates used in this study, the line focus at the BFP extends along the *x* axis, where the length of the line at the BFP determines the magnitude of NA_*x*_. For SPIM, a relatively low NA_*x*_ is used to generate a light sheet that maintains its thickness over a relatively long axial propagation distance (i.e., long Rayleigh range) but at the expense of a thicker sheet (larger beam waist). Since the line focus at the BFP is narrow along the *y* axis (NA_*y*_ ~ 0), there is no beam focusing in the *y* direction (i.e., the beam is collimated in the *y* direction). While simple and straightforward, this illumination method has two main drawbacks. First, the lack of angular diversity in the *y*-direction, and minimal angular diversity in the *x* direction (for a low-NA light sheet), results in illumination shadowing artifacts within the sample due to occlusions [17]. Second, the illumination light sheet is scattered (Mie and Rayleigh scattering) in biological tissues, which generates an unwanted background that reduces image contrast (signal-to-background ratio, SBR), and consequently, imaging depth.

In the mSPIM design (Fig. 1c), a cylindrical lens is used to focus a line onto a pivoting mirror positioned at a conjugate front focal plane (FFP*) of the illumination objective. The pivoting mirror, combined with a tube lens, effectively translates the line focus at the BFP of the illumination objective (in the *y* direction), resulting in a rotation of the 2D light sheet within the sample over time (*x*-axis rotation), which provides sufficient angular diversity (if a time-averaged image is obtained) to mitigate the shadows cast by occluding objects (Fig. 1d). When compared to SPIM, NA_*x*_ remains unchanged, whereas the effective NA_*y*_ is enlarged by a given rotation angle (in a time-averaged sense). However, like SPIM, mSPIM still utilizes a 2D light sheet with widefield camera detection, which can lead to poor image contrast. In addition, the imaging framerate (i.e., speed) is constrained by the time it takes to physically rotate the light sheet through at least one full range (i.e. half a sinusoidal period) per frame.

To enhance image contrast with LSFM, digital scanned light-sheet microscopy (DSLM) with confocal line detection was developed [5, 19, 20]. Unlike SPIM and mSPIM, DSLM utilizes a circular Gaussian pencil beam with lateral symmetry (NA_*x*_ = NA_*y*_), which is focused onto a scanning mirror positioned at a conjugate BFP (Fourier plane) of the illumination objective (Fig. 1e). The scanning mirror, combined with a scan and tube lens, translates the pencil beam along the *y* axis within the sample, generating a 2D light sheet over time. The position of the scanned pencil beam can be synced to the rolling shutter of a detector array, which acts as a digital confocal slit. This approach, commonly referred to as confocal line detection, rejects out-of-focus scattered light with a slit whose thickness, *ω*_*siit*_, approximates the beam-waist diameter of the pencil beam, 2*ω*_0,*y*_, yielding enhanced image contrast. However, to achieve the long Rayleigh range that is typically desired for the pencil beam (and the resultant digitally scanned 2D light sheet), a relatively low NA is typically used, resulting in minimal angular diversity in the *y* direction and similar (though slightly reduced) shadowing artifacts as SPIM.

The multidirectional digital scanned light-sheet microscopy (mDSLM) architecture utilizes an elliptical Gaussian pencil beam with a higher NA along one axis (in the plane of the light sheet) to provide increased angular diversity for mitigation of shadowing artifacts, and a lower NA along the axis orthogonal to the light sheet in order to maintain a long depth of focus (i.e. a light sheet that maintains its thickness over a relatively long propagation distance). From Eq. (1), it is apparent that the beam radii, waists, and Rayleigh ranges given by Eqs. (2-4) are independent, and that an elliptical Gaussian beam (i.e., a beam with a different NA in the *x* and *y* directions) can be utilized to combine the benefits of the mSPIM and DSLM architectures. Unlike mSPIM, where the angular diversity in the *y* direction is achieved by rotating a 2D light sheet over time, mDSLM utilizes a passive elliptical Gaussian beam that provides similar angular diversity in the *y* direction (without rotation), and therefore does not impose additional constraints on speed.

**Figure 1.**
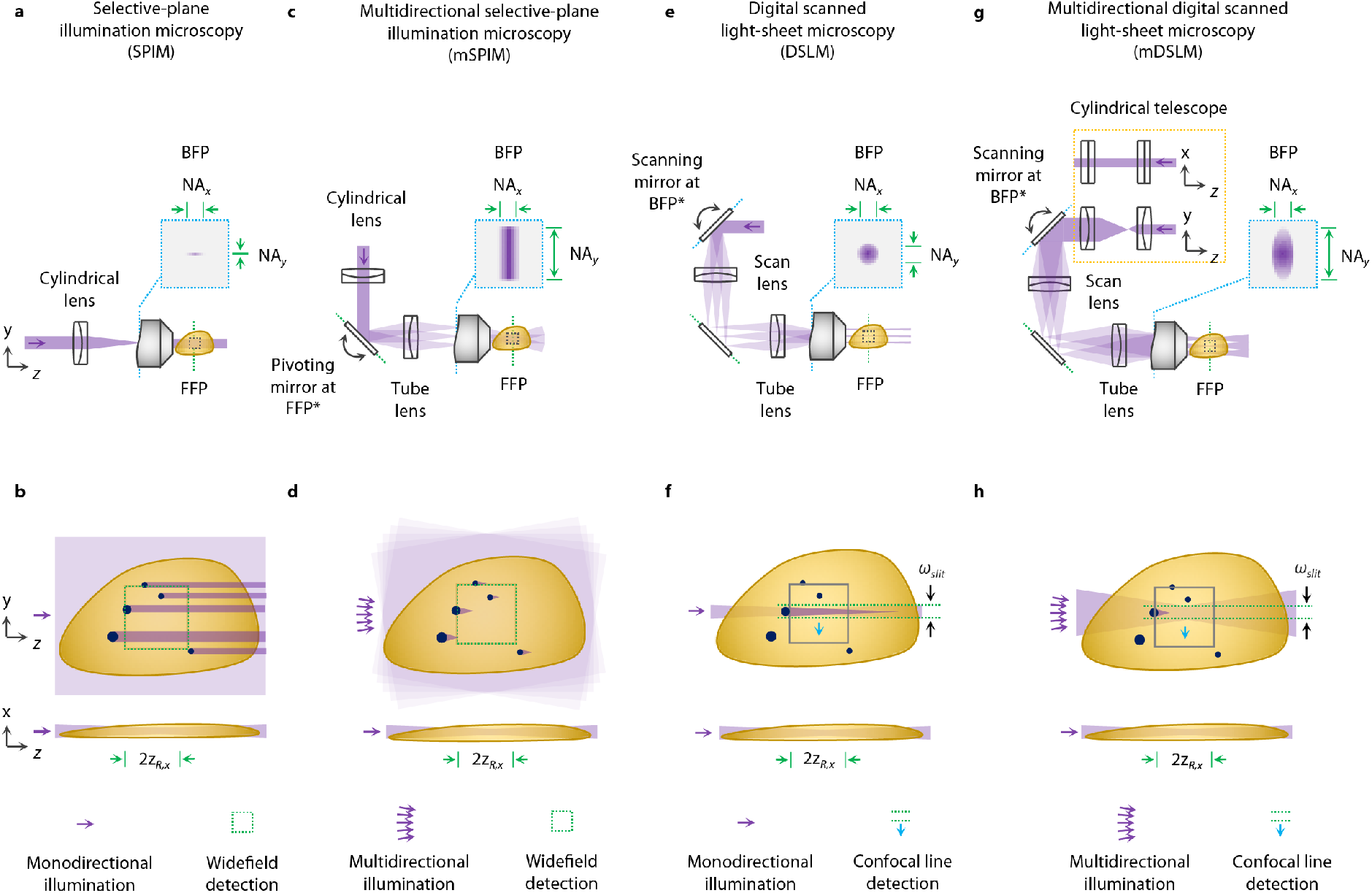
Comparison of SPIM, mSPIM, DSLM, and mDSLM architectures. (a) The optical layout of selective plane illumination microscopy (SPIM) is shown [9], A cylindrical lens is used to create a focal line at the back focal plane (BFP) of an illumination objective, generating a static 2D light sheet within the sample, (b) A zoomed-in view of the illumination path within the sample. Due to a lack of angular diversity in the 2D light sheet, strong shadowing artifacts are visible due to occlusions. Widefield camera detection is used (dashed green box in panel b) to image the static 2D light sheet. (c) Multidirectional selective plane illumination microscopy (mSPIM) uses a pivoting mirror, located at a conjugate front focal plane (FFP*), to translate the line focus across the BFP of the illumination objective [18]. (d) This results in a 2D light sheet that rotates within the sample (in the *yz* plane) to average out the shadowing artifacts over time. (e) Digital scanned light-sheet microscopy (DSLM) uses a circular Gaussian pencil beam at the BFP of the illumination objective, and a scanning mirror located at a conjugate back focal plane (BFP*), to translate the pencil beam within the sample in the *y* direction, creating a 2D light sheet over time [5]. (f) To achieve a long Rayleigh range (a long depth of focus) a relatively low NA is used, resulting in minimal angular diversity in the *x* and *y* directions and similar (though slightly reduced) shadowing artifacts as SPIM. As the pencil beam in DSLM is scanned in *y*, it is synchronized with the rolling shutter of a sCMOS camera (dashed green lines), which acts as a confocal slit to reject background light and improve image contrast [19, 20]. (g) Multidirectional digital scanned light-sheet microscopy (mDSLM) is similar to DSLM, with the addition of a cylindrical telescope to generate an elliptical Gaussian beam at the BFP of the illumination objective such that NA*y* is increased while NA*x* is unaltered. This results in a long Rayleigh range in the *x* direction (a light sheet that maintains its thickness over a relatively long propagation distance), but with increased angular diversity in the *y* direction to mitigate shadowing artifacts. The illumination and detection characteristics of the SPIM, mSPIM, DSLM, and mDSLM architectures are symbolically depicted at the bottom of panels (b), (d), (f), and (h).

### Illumination around large refractive objects and optimization of beam parameters (DSLM vs. mDLSM)

A large glass sphere (diameter, *d* = 20 μm and refractive index *n*_*sphere*_ = 1.59) was embedded within a fluorescent gel with a refractive index *n*_*gel*_ = 1.46. The sphere was positioned at a depth of *z*_*sphere*_ = 125 μm at an offset of Δ*y* = 2 μm from the optical axis of the pencil beam. The pencil beam focus was located at a depth of *Z*_*focus*_ = 350 μm. The propagation characteristics of both the DSLM and mDSLM beams around the glass sphere were simulated and measured experimentally from a starting position of *z*_*0*_ = 0 μm to *z* = 700 μm.

**Figure 2.**
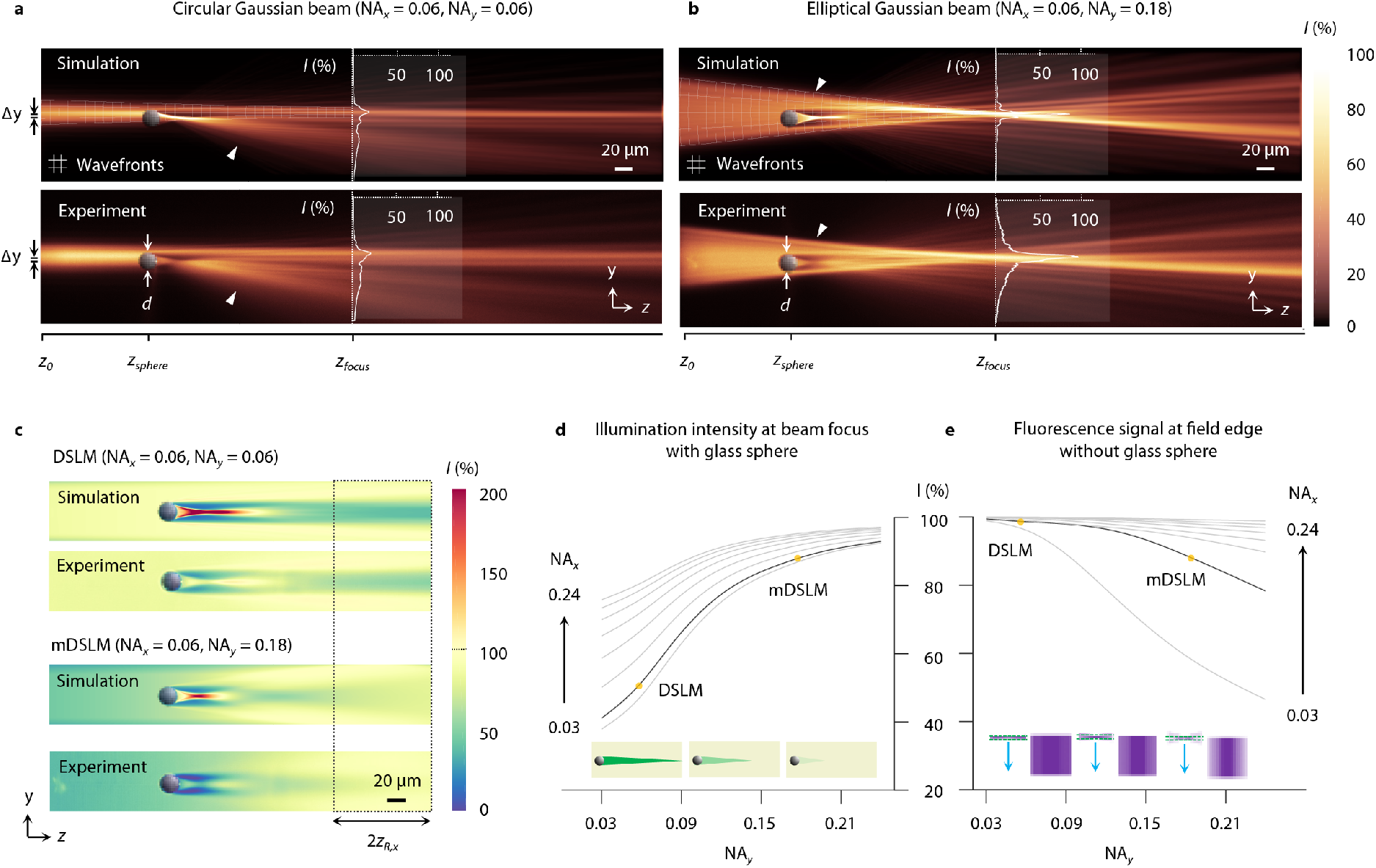
Illumination around large refractive objects and optimization of beam parameters (DSLM vs. mDLSM). (a) Simulation and corresponding experimental image of a circular Gaussian beam (NA_*x*_ = NA_*y*_ = 0.06) propagating around a large glass sphere (diameter *d* = 20 μm, *n*_*sphere*_ = 1.59) embedded within a fluorescent gel (*n*_*gel*_ = 1.46). The sphere is positioned at a depth of *z*_*sphere*_ =125 μm at an offset of Δ*y* = 2 μm from the optical axis of the pencil beam. The pencil beam focus is located at a depth of *Z*_*focus*_ = 350 μm. For a circular Gaussian beam (DSLM), the intensity at the beam focus is reduced by >75% relative to an unobstructed beam, as illustrated by the overlaid line profiles. (b) Simulation and corresponding experimental image of an elliptical Gaussian beam (NA_*x*_ = 0.06, NA_*y*_ = 0.18) propagating through an identical fluorescent gel and glass sphere. In contrast to the circular Gaussian beam used in DSLM, the increased angular diversity in the *y* direction enables the elliptical Gaussian beam (used in mDLSM) to experience only a ~10% reduction in intensity relative to an unobstructed beam. Simulated and experimental beam-scanned images, with DSLM and mDLSM, are shown in (c). Simulation results are plotted in (d) for the dependence of the beam-focus intensity (behind the glass sphere) as a function of NA (NA_*x*_ = 0.03 - 0.24 and NA_*y*_ = 0 - 0.24, in increments of 0.03). Panel (e) provides simulation results for the dependence of the fluorescence signal at the field edge, as a function of NA_*x*_ and NA_*y*_ (in the absence of the glass sphere). In both (d) and (e), the black solid line indicates the value of NA_*x*_ (0.06) used experimentally in the majority of our studies, in which the experimentally used values of NA_*y*_ are indicated as yellow points (0.06 and 0.18 for DSLM and mDSLM, respectively). Illustrations are shown above the horizontal graph axes in (d) and (e) to depict the reduction in shadowing artifact as a function of NA_*y*_ (d), but an increasing roll off in signal at the field edge (e).

For light propagation around a glass sphere, the intensity, *I*, for a circular and elliptical Gaussian illumination beam were calculated using a recently published simulation method based upon the beam propagation method (BPM) [30]. For the circular Gaussian beam used in DSLM, NA_*x*_ = NA_*y*_ = 0.06, whereas for the elliptical Gaussian beam used in mDSLM, NA_*x*_ = 0.06 and NA_*y*_ = 0.18. The results are shown in Figs. 2a and 2b.

The simulation results show that the standard DSLM beam is severely occluded by the glass sphere, resulting in >75% reduction in intensity at the beam focus relative to the intensity distribution in the absence of a glass sphere. In comparison, the mDLSM beam only experiences a ~10% reduction in intensity at the beam focus due to the glass sphere. The difference in the angular diversity of both beams through the glass sphere is depicted by the overlaid wavefront grids shown in the *yz* plane. In the corresponding experimental measurements, shown below the simulations, the same trends are observed. Both simulated and experimentally recorded illumination intensities were normalized to the intensity at the beam focus in the absence of a glass sphere.

To investigate the shadowing artifacts that would be observed during DSLM and mDSLM imaging, simulations and experiments were performed with laterally scanned beams, in which confocal line detection was used. Similar to a previous study [20], the slit size,*ω*_*slit*_, was chosen to be 1.5× the beam diameter at the Rayleigh range 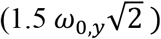. Note that although the mDLSM beam has a higher NA_*y*_ than the DSLM beam, an identical slit size was used (based on the DSLM beam) to provide a similar degree of background rejection and optical sectioning. The results in Fig. 2c reveal that in both simulations and experiments, the intensity at the light sheet focus for DSLM is reduced by ~50%, compared to ~10% for mDLSM. The usable field of view in the x direction is also indicated in Fig. 2c, corresponding to a confocal parameter (depth of focus) of 2*z*_*R,x*_ ~ 100 μm. A video comparing the simulated and experimentally measured propagation of DSLM (circular) and mDLSM (elliptical) beams around the glass sphere are shown in Supplementary Video 1.

To further explore the dependence of the shadowing artifacts on both NA_*x*_ and NA_*y*_, numerical simulations similar to the results shown in Fig. 2c were calculated for NA_*x*_ = 0.03 - 0. 24 and NA_*y*_ = 0 - 0.24, both in increments of 0.03. The intensities at the beam focus are plotted in Fig. 2d. In general, increasing the angular diversity by maximizing NA_*y*_ causes the intensity at the beam focus to remain high (i.e. it reduces shadowing artifacts). However, there is a decreasing benefit to increasing NA_*y*_ as NA_*x*_ is increased. This is due to the fact that as NA_*x*_ is increased, it introduces sufficient angular diversity such that utilizing an elliptical Gaussian beam (with a much larger NA_*y*_) is no longer needed for reducing shadowing artifacts. Note that for most LSFM systems, a relatively low NA_*x*_ is desired to generate a long depth of focus in which the light sheet thickness is relatively constant over a long axial extent (propagation distance). Finally, there appears to be a marginal benefit to increasing NA_*y*_ beyond ~0.20, which corresponds to a focusing angle of ±10 deg. This is consistent with the pivoting angle typically used for mSPIM [18].

One consequence of the elliptical Gaussian beam used in mDSLM is a decrease in the fluorescence signal at the edges of the field of view (i.e. at ±*z*_*R,x*_) as NA_*y*_ is increased. This is due to the more-rapid expansion of the Gaussian pencil beam in the *y* direction as one moves away from the beam waist (for a higher-NA beam), which causes the beam to overfill the confocal slit. To explore this tradeoff, simulations over a range of NA_*x*_ and NA_*y*_ (same range as previously explored) were conducted in the absence of the glass sphere, in which the signal was recorded at the field edge (i.e. at ±*z*_*R,x*_). The results are plotted in Fig. 2e, showing that as NA_*y*_ is increased, the signal at the beam edge is decreased. An alternative would be to increase the size of the confocal slit for an mDSLM system in order to minimize signal loss at the field edges, but at the expense of reduced background rejection and degraded contrast. Therefore, there is a balance in selecting NA_*y*_ for a given NA_*x*_ to optimize a mDSLM system, and in choosing an optimal confocal slit size. The focusing parameters used in the majority of this study for both DSLM (NA_*x*_ = NA_*y*_ = 0.06) and mDSLM (NA_*x*_ = 0.06, NA_*y*_ = 0.18) are indicated in Figs. 2d and 2e. Note that in Fig. 2e, the signal roll-off at the field edge is negligible for DSLM, and is ~12% for mDSLM.

### Imaging through small refractive heterogeneities (DSLM vs. mDSLM)

To further demonstrate the ability of mDSLM to mitigate shadowing artifacts in comparison to DSLM, experiments was performed with fluorescent gels (*n*_*gel*_ = 1.46) containing a multitude of small glass spheres of diameter, *d* = 6 μm, and refractive index, *n*_*sphere*_ = 1.59. For DSLM, NA_*x*_ = NA_*y*_ = 0.06, and for mDSLM, NA_*x*_ = 0.06 and NA_*y*_ = 0.18, resulting in a matched depth of focus in the *x* direction of 2*z*_*R,x*_ ~ 100 μm for both imaging methods, within which the light sheet thickness remains relatively constant. DSLM and mDSLM images were acquired in 50-μm steps along the *z* axis, analogous to tiling SPIM [31]. In order to quantify the changes in illumination intensity, Δ*I*, all images were normalized by images recorded in an identical homogeneous fluorescent gel containing no glass spheres. The results are shown in Figs. 3a and 3b.

**Figure 3.**
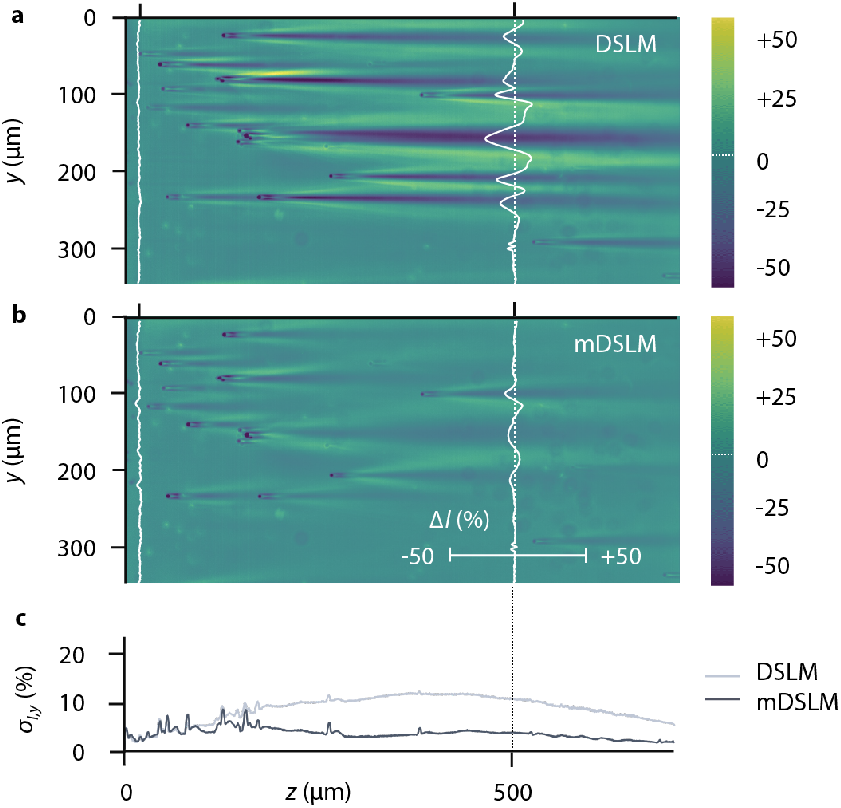
Imaging through small refractive heterogeneities (DSLM vs. mDSLM). (a) An image generated by DSLM (NA_*x*_ = NA_*y*_ = 0.06) through a cluster of glass spheres. Shadowing artifacts, with intensity deviations on the order of ±30%, are visible behind each glass sphere. A corresponding mDSLM (NA_*x*_ = 0.06, NA_*y*_ = 0.18) image is shown in (b). Due to increased angular diversity in the *y* direction, the intensity deviations caused by shadowing artifacts are reduced to ±12%. Line profiles at a depth of *z* = 500 μm are shown. In (c), the standard deviation of the intensity fluctuations along the *y* axis, σ_*I,y*_, is plotted as a function of *z* for the DSLM and mDSLM images. At all depths beyond *z* = 100 μm, σ_*I,y*_ is greater for DSLM compared to mDSLM.

For DSLM, large streaks and shadows due to the glass spheres are visible, causing intensity deviations on the order of ±30%. In comparison, images of the same phantom using mDSLM exhibit reduced streaks and shadows with intensity deviations on the order of ±12%. Line profiles through the recorded images at a depth of *z* = 500 μm are shown. To quantify the severity of the shadowing artifacts for DSLM and mDSLM, the standard deviation in the illumination intensity along the *y*-axis, *σ*_*I,y*_, for each *z* position, was calculated. The results, plotted in Fig. 3c, show that the standard deviation for DSLM is as much as 3× higher than for mDSLM due to the accumulation of more intense and persistent streaks and shadows.

### mDSLM enhances imaging contrast in comparison to mSPIM

Experiments were conducted to explore the differences in image contrast between mDSLM, which utilizes confocal line detection, and mSPIM, which uses widefield detection. In these experiments, non-fluorescent gels (*n*_*gel*_ = 1.46) with fluorescent glass spheres (*n*_*sphere*_ = 1.59) of diameter *d* = 7 μm were imaged, where tissue scattering was generated by mixing sub-micron lipid droplets (Intralipid) in the non-fluorescent gel at a volume concentration of ~1% and refractive index mismatch of Δ*n* = *n*_*Upid*_ − *n*_*gel*_ ~ 0.1 [32]. This level of refractive index mismatch is comparable to that of optically cleared tissues that are commonly imaged using LSFM [33, 34]. For mDSLM, NA_*x*_ = 0.06, NA_*y*_ = 0.18, and for mSPIM, NA_*x*_ = 0.06 with a pivoting angle of ~10 deg., which is equivalent to NA_*y*_ ~ 0.18 (after time averaging). For both imaging methods, the depth of focus in *x* was 2*z*_*R,x*_ ~ 100 μm, and images were acquired by tiling in 50 μm steps along the *z*-axis to a depth of 1200 μm.

**Figure 4.**
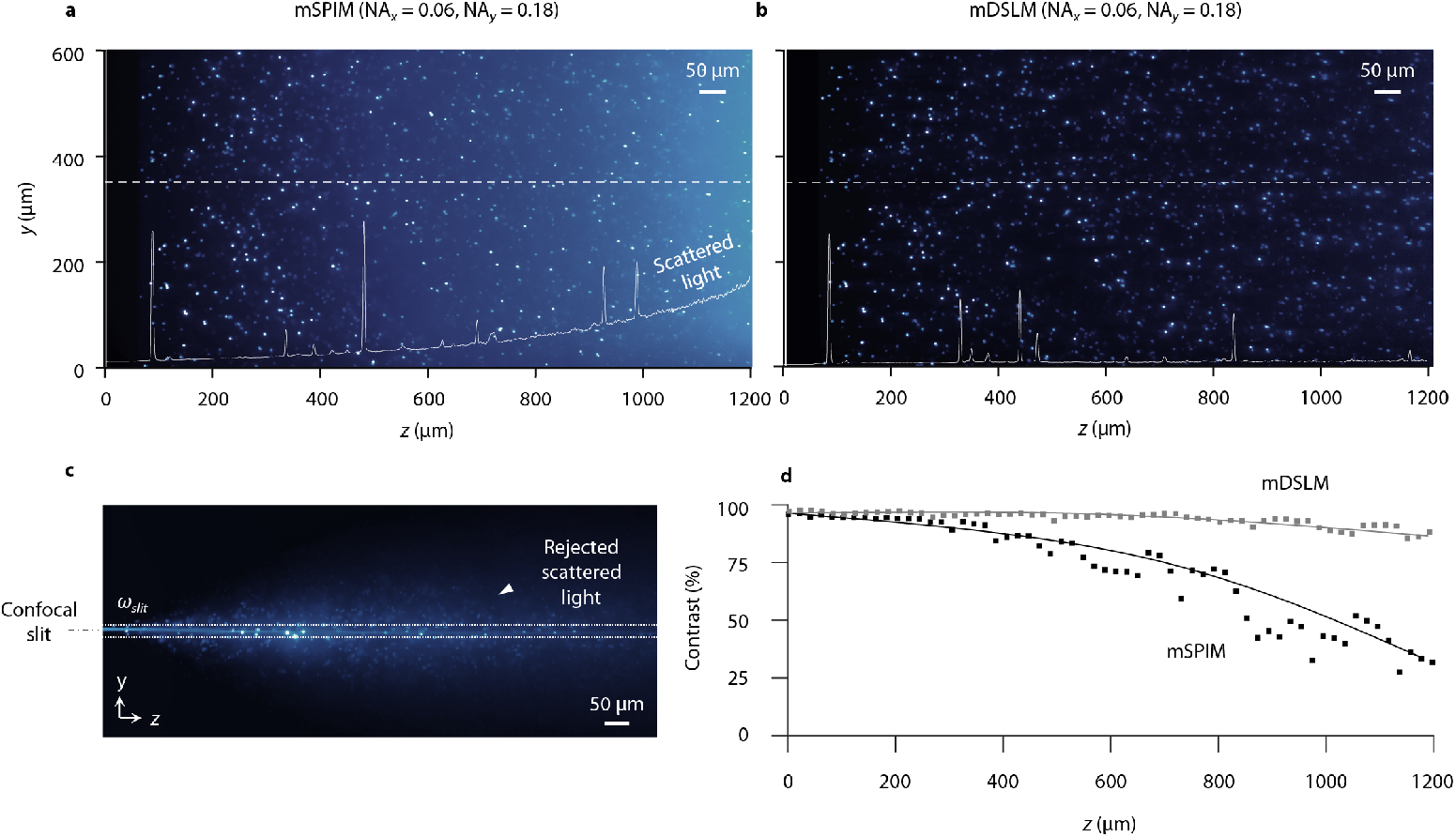
mDSLM enhances imaging contrast in comparison to mSPIM. (a) An image generated by mSPIM (NA_*x*_ = 0.06, NA_*y*_ = 0.18) in a non-fluorescent gel containing fluorescent glass spheres (diameter = 7 μm). A 1% volume concentration of sub-micron lipid droplets was mixed into the gel to simulate a small amount of tissue scattering. mSPIM uses widefield camera detection and is therefore unable to reject the out-of-focus scattering background. This is visible in the overlaid line profile at *y* = 350 μm, which shows an increasing scattering-induced background as a function of *z*. The corresponding mDSLM (NA_*x*_ = 0.06, NA_*y*_ = 0.18) image is shown in (b). By using a laterally scanned elliptical Gaussian pencil beam to generate a 2D light sheet, mDSLM is compatible with confocal line detection, which rejects much of the scattering background. An image of a stationary Gaussian pencil beam (at *y* = 0 μm) is shown in (c), in which the size of the confocal slit is displayed. An example of out-of-focus background light due to tissue scattering is highlighted by the inset arrow. Image contrast as a function of *z* is plotted in (d), at 20 μm intervals, for both mSPIM and mDSLM. At *z* = 1200 μm, the contrast for mSPIM is reduced to ~30%, whereas for mDSLM the contrast remains >90%.

For mSPIM, the use of widefield camera detection leads to the increased collection of background scattered light at deeper depths, and therefore reduced imaging contrast (Figs. 4a and 4b). On the other hand, mDSLM is able to reject out-of-focus portions of the scattered light in the *y* direction by using confocal line detection. These findings are reinforced by the line profiles through the images at *y* = 350 μm. A representative image of a stationary Gaussian pencil beam used for mDSLM is shown in Fig. 3c. The size of the confocal slit, ω_*siit*_, is shown, where the inset arrow highlights the out-of-focus scattered light that is rejected by the slit. It should be noted that the confocal slit is only effective at rejecting out-of-focus light along the *y* direction and not the *x* direction. All images were corrected for the exponential attenuation of the illumination light as a function of depth to yield a normalized maximum image intensity as a function of *z* (see Methods).

To quantify the contrast enhancement with mDSLM relative to mSPIM, the image contrast, *C* = (*I*_*max*_ − *I*_*min*_)/(*I*_*max*_ + *I*_*min*_), was calculated as a function of *z* for 20-μm wide regions of interest (Fig. 4d). To a depth of 1200 μm, mDSLM maintains an image contrast > 90%, whereas for mSPIM the image contrast degrades to ~30%.

### mDSLM mitigates shadowing artifacts and enables contrast-enhanced imaging in biological tissues

The imaging performance of mDSLM in comparison to SPIM, mSPIM, and DSLM was assessed in fluorescently labeled biological tissues, which exhibit a wide distribution of scattering properties, occlusions, and refractive heterogeneities [35, 36].

In a first set of experiments, optically cleared human breast tissue was labeled with eosin. The intricate combination of adipose and stroma in human breast tissue represents a biological structure similar to that of glass spheres in a fluorescent gel phantom. 2D images were generated through tiled acquisition at 50-μm increments along the *z*-axis to a depth of 1000 μm. The results are shown in Figs. 5a - 5c. For SPIM, both shadowing artifacts and reduced imaging contrast are observed due to the lack of angular diversity in the light sheet as well as the use of widefield detection (no confocal slit). mSPIM mitigates the shadowing artifacts but still shows reduced image contrast. On the other hand, DSLM provides enhanced image contrast but still generates shadowing artifacts. By combining passive multidirectional illumination with confocal line detection, mDSLM enables both mitigation of shadowing artifacts and enhanced image contrast. Zoomed-in views of DSLM versus mDSLM, and mSPIM versus mSPIM, are shown in Fig. 5b, with intensity line profiles plotted in Fig. 5c. Additional comparison images of human breast tissue are shown in Supplementary Figures 1 and 2.

In a second set of experiments, the small intestine of a mouse was optically cleared and labeled with acridine orange (primarily a nuclear stain). The mucosa of the small intestine provides a layered glandular structure for comparing the four LSFM methods both in the vertical (depth-wise) and *en face* planes (parallel to the tissue surface). Three-dimensional (3D) images were acquired by tiling along the *z*-axis at 50-μm increments to a depth of 600 μm, as well as stage-scanning the sample at a sampling pitch of 0.55 μm per pixel in the *x* direction over a distance of 300 μm. The results are shown in Figs. 5d - 5f. The *en face* images in the yz plane (Figs. 5b and 5e) show similar trends to the vertical images in Figs. 5a and 5d. The images shown in Fig. 5e are at *z* positions (depths) of 100, 250, 400, and 550 μm. At superficial depths, the crypts and villi of the small intestine are largely free of aberrations and artifacts for all four LSFM methods. However, at deeper *z* positions, the shadowing artifacts appear as striated dark and bright patches in the SPIM and DSLM images. In addition, the SPIM and mSPIM images exhibit reduced image contrast at greater depths. Line profiles through the intestinal crypts at *z* = 550 μm are plotted in Fig. 5f, along with corresponding values for contrast, *C* = (*I*_*max*_ − *I*_*min*_)/(*I*_*max*_ + *I*_*min*_). All images were corrected for the exponential attenuation of the illumination light to yield a normalized maximum image intensity as a function of *z* (see Methods section). Videos comparing the image quality of the *z* stacks acquired using SPIM, mSPIM, DSLM, and mDSLM are shown in Supplementary Video 2.

**Figure 5.**
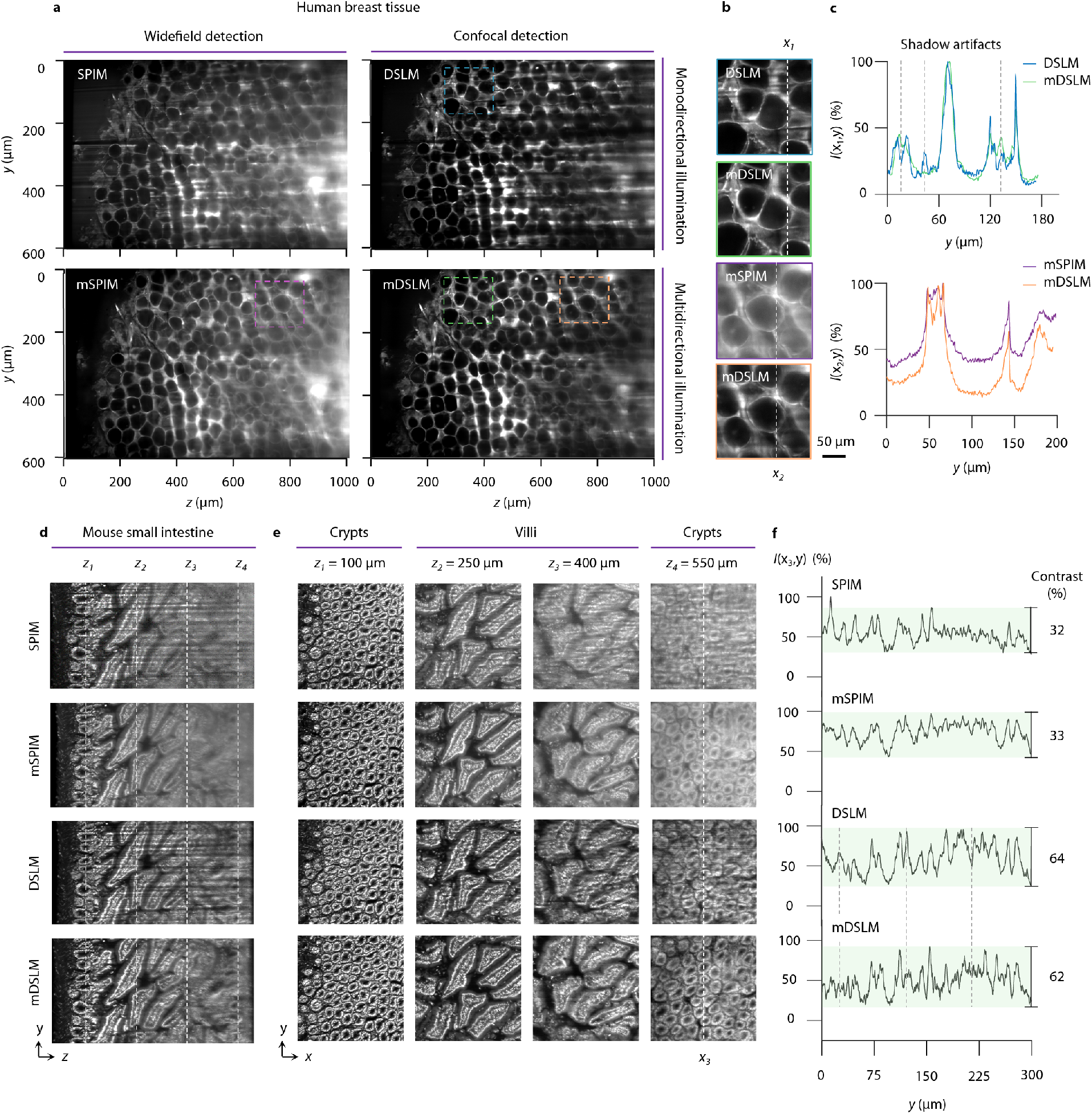
mDSLM mitigates shadowing artifacts and enables contrast-enhanced imaging in biological tissues. SPIM, mSPIM, DSLM, and mDSLM images of human breast tissue stained with eosin are shown in (a). Shadowing artifacts are visible in the SPIM and DSLM images, whereas reduced contrast is observed in the SPIM and mSPIM images. Representative zoomed-in views demonstrating the mitigation of shadowing artifacts for mDSLM versus DSLM, and enhanced imaging contrast for mDSLM versus mSPIM, are shown in (b). Intensity line profiles through the zoomed-in views in (b) are plotted in (c). The dashed lines in (c) mark the *y* positions of the shadows seen in the DSLM image. SPIM, mSPIM, DSLM, and mDSLM images of murine small intestine, stained with acridine orange, are shown in (d). *En face* images in the *xy* plane are shown in (e) at *z* positions (depths) of 100, 250, 400, and 550 μm. Intensity line profiles along the dashed line in the images of the crypts at *z* = 550 μm are plotted in (f), with values for image contrast shown on the right. The dashed lines in (f) mark the *y* positions of the shadows seen in the DSLM images in (d) and (e), and which are mitigated in the mDSLM images.

## Discussion

In recent years, LSFM has become a powerful imaging tool for a variety of biological investigations, and has also shown promise for applications in clinical pathology [2–12, 14, 15, 29]. Although LSFM is conventionally used to image highly transparent objects such as embryos, single cells, and optically cleared tissues, residual artifacts still exist due to imperfect homogenization of the refractive-index distribution in biological specimens. For example, shadowing artifacts and scattering-induced background light both often contribute to poor image quality [17], which can lead to erroneous biological findings and inaccurate clinical determinations. The presence of these artifacts in the original SPIM architecture, which utilizes a static 2D light sheet, spurred the development of mSPIM, a technique for pivoting a 2D light sheet to average out shadows over time, as well as the development of DSLM, which enhances image contrast by scanning a pencil beam that is synchronized to a confocal rolling shutter to reject out-of-focus and multiply scattered light [5, 18–20]. Unfortunately, the mSPIM and DSLM approaches are not compatible since rotating the pencil beam used in DSLM would cause much of the beam to rotate out of the confocal slit. In addition, the rotation would have to be approximately two orders of magnitude faster than what is necessary for mSPIM in order to match the integration time of a rolling shutte (see Supplementary Materials for additional details).

Here we have demonstrated a multidirectional DSLM technique (mDSLM), that utilizes an elliptical Gaussian pencil beam in which the numerical apertures are decoupled in the directions parallel and orthogonal to the light sheet (NA_*x*_ ≠ NA_*y*_). A low NA in the direction orthogonal to the light sheet (NA_*x*_) is used to maintain a long Rayleigh range in *x* (i.e. a light sheet that maintains its thickness over a relatively long axial propagation distance) while a higher NA in the plane of the light sheet (NA_*y*_) is used to generate angular diversity for the mitigation of shadowing artifacts. With mDLSM, increased angular diversity is passively provided in the beam itself, rather than generated by physically pivoting a beam over time (as with mSPIM). As a result, confocal line detection is possible (as with DSLM), as was described in the introduction, without additional speed constraints.

While we chose not to focus on it in this study, an additional passive multidirectional illumination approach, which is compatible with both SPIM and DSLM, is to utilize a diffraction grating positioned at a conjugate front focal plane (FFP*) of the illumination objective to increase angular diversity in *y* (see Supplementary Figures 4 - 6) [37]. A grating-based approach directly generates angled 2D light sheets or 1D pencil beams at discrete angles within the sample. However, this approach is inefficient (typically there is power loss through transmission diffraction gratings), not achromatic (it is difficult to engineer a transmission diffraction grating which splits several incident wavelengths at the same angle), only increases the NA_*y*_ at discrete angles, and results in an undesirable interference pattern due to the coherence of the various angled light sheets and pencil beams that overlap within the sample. This interference pattern must be time-averaged away by slightly dithering (spatially translating) the interference pattern within each camera exposure. Despite these drawbacks, the use of a diffraction grating to generate multiple angled 2D light sheets has a significant speed advantage over traditional mSPIM. While mSPIM requires the pivoting mirror to rotate a light sheet over its full angular range (~10 deg) within the framerate of the imaging camera, the use of a diffraction grating only requires the illumination beam to be rotated or translated enough to cause peaks in the interference pattern to move to the location of adjacent peaks (see Supplementary Figures 7 and 8). As a result, within a single scanning period (e.g. with a pivoting mirror), multiple shadow-free images can be acquired (~100 images, as shown in Supplementary Figure 8), reducing the speed requirements of the scanning mirror by several orders of magnitude. Videos and figures comparing the simulated and experimentally measured propagation of mSPIM and mDSLM beams generated with a diffraction grating are shown in Supplementary Videos 3 and 4 and Supplementary Figure 9.

Another alternative for reducing shadowing artifacts is the use of a propagation invariant beam, such as a Bessel or Airy beam [14, 26–28]. However, while such beams do exhibit “self-healing” properties, and therefore mitigation of shadowing artifacts, the out-of-focus side lobes that are necessary for self-healing also result in reduced image contrast, even when combined with confocal line detection. While advancements in computational deconvolution algorithms and technologies are currently in development (including the use of graphics processing units), LSFM datasets are notoriously large, often terabytes in size, which can make an analog approach attractive for minimizing shadowing artifacts and maximizing image contrast [23].

In summary, the mDSLM approach is advantageous in that it is a simple and passive method that does not rely on post-processing and can be readily incorporated into a standard DSLM architecture by inserting a cylindrical telescope to expand the NA along one axis. The mDSLM approach mitigates shadowing artifacts and is compatible with confocal line detection for contrast-enhanced imaging without imposing additional constraints on speed. More generally, the mDLSM approach demonstrates that decoupling the NA of the illumination beam along two orthogonal axes can provide an additional degree of freedom for the design and optimization of LSFM systems. Ultimately, the ability to rapidly generate 3D microscopy datasets with high imaging fidelity and optimal contrast/depth, as enabled by mDLSM, will be of value for ensuring accurate biological observations and clinical determinations.

## Methods

### Optical setup and image acquisition

A custom LSFM system was used for all experiments (see Supplementary Figure 1). Light from a 0.12 NA fiber-coupled laser (488 or 660 nm) was collimated by a lens L1 (*f* = 19 mm) (AC127-090, Thorlabs). Light was then directed to a beam-shaping module that enabled rapid switching between the SPIM, mSPIM, DSLM, and mDSLM architectures (see Supplementary Figures 10-13 for the experimental setups). For SPIM and mSPIM, collimated light was focused to a line at a conjugate back focal plane (BFP*) of the illumination objective using a cylindrical lens, C1 (*f* = 50 mm) (ACY254-050, Thorlabs). For DSLM, the cylindrical lens was removed, and a standard circular Gaussian beam was relayed to the BFP* of the illumination objective. Finally, for mDSLM, a 3× cylindrical telescope comprised of two cylindrical lens, C1 (*f* = 50 mm) (ACY254-050, Thorlabs), C2 (*f* = 150 mm) (ACY254-150, Thorlabs) was used to elongate the Gaussian beam in the *y*-direction at the BFP*.

Light was then relayed by two lenses, L3 (*f* = 50 mm) (AC254-075, Thorlabs), L4 (*f* = 75 mm) (AC254-075, Thorlabs) and focused onto a pivoting mirror (6210H, Cambridge Technology) positioned in a conjugate front focal plane (FFP*) of the illumination objective by a third lens, L5 (*f* = 200 mm) (AC254-200, Thorlabs). Light from the mirror was collected by a scan lens L6 (*f* = 70 mm) (CLS-SL, Thorlabs) and focused onto a scanning mirror (6210H, Cambridge Technology) positioned in a second BFP* of the illumination objective. Finally, light from the second scanning mirror was collected by a second scan lens L6 (*f* = 70 mm) (CLS-SL, Thorlabs) and imaged onto the BFP of the illumination objective (XLFLUOR/340 4X, 0.28 NA, Olympus) using a tube lens L7 (*f* = 165 mm) (TTL-165, Thorlabs).

For SPIM, both the pivoting and scanning mirrors were turned off. For mSPIM, the pivoting mirror in the FFP* was actuated to rotate the beam within the sample. For DSLM and mDSLM, the pivoting mirror was turned off, and the scanning mirror located in the BFP* was actuated to scan the beam within the sample. Both mirrors were driven using a function generator with a 50-Hz sawtooth voltage (DS345, Stanford Research Systems). The voltage amplitude of the sawtooth was experimentally calibrated to either provide a pivoting angle of ~10 deg within the sample (for mSPIM) or to scan across the full field of view of the collection objective (for DSLM and mDSLM). The illumination optics and scanning mirrors, combined with the BFP diameter and 0.28 NA of the illumination objective, yield an effective NA_*x*_ ~ 0.06, NA_*y*_ = 0 (for SPIM); NA_*x*_ = NA_*y*_ ~ 0.06 (for DSLM); NA_*x*_ ~ 0.06, NA_*y*_ ~ 0.18 (for mSPIM); and NA_*x*_ ~ 0.06, NA_*y*_ ~ 0.18 (for mDSLM).

The illumination light was transmitted through a fused quartz glass cuvette (*n* = 1.46) that contained samples immersed in a refractive index-matching solution (*n* = 1.46). The fluorescence excited within the samples was collected using an objective (Cleared Tissue Objective, 16.3X, 0.40 NA, Applied Scientific Imaging/Special Optics), transmitted through a bandpass fluorescence filter, and imaged onto a sCMOS camera (ORCA Flash 4.0 v2, Hamamatsu) using a tube lens, L8 (*f* = 100 mm) (TTL-100, Thorlabs). For all experimental images, the focal plane of the imaging objective was positioned approximately 100-μm deep within the agarose phantom and biological specimens. The chosen collection optics resulted in a sampling pitch of ~0.77 μm/pixel.

Tiled 2D and 3D datasets were collected using a custom LABVIEW program (2016, 64- bit, National Instruments) and post-processed with a combination of MATLAB (R2017, Mathworks) and ImageJ running on a local desktop workstation (Windows, 64-bit, 256 GB RAM, 3.7 GHz processor, 8 TB RAID0 HD array, TitanXP GPU). For all experimental measurements, the camera was operated at 50 frames per second (exposure time of 50 ms per pixel for widefield imaging and ~200 μs per pixel for confocal line detection with *ω*_*slit*_ = 20 μm or 25 pixels).

### Numericai simuiations

Numerical simulations were executed on the local workstation using a previously described BPM simulation architecture in MATLAB (R2017, Mathworks) [30]. For the results shown in Fig. 2, the 3D illumination intensity distribution, *I*(*x*,*y*,*z*), was recorded in a 300 by 300 by 700 μm (*xyz*) volume with a voxel size of 0.25 μm in all three dimensions. The refractive index of the entire medium, *n*(*x*,*y*,*z*), was set to *n*_*gel*_ = 1.46, except for the glass spheres which were set to *n*_*sphere*_ = 1.59. The computed 3D illumination intensity distribution was multiplied by fluorescence values in each voxel, *F*(*x*,*y*,*z*), (*F* = 1 for every voxel except within the voxels occupied by the glass sphere, where *F* = 0), and convolved with the orthogonal 3D point spread function of the collection objective, *C*(*x*,*y*,*z*) (simulated for the 0.40 detection NA used experimentally) to obtain a final fluorescence image simulation, *S* = (*I*×*F*)⨂*C*.

To simulate scanning, the beams were scanned in 0.25 μm increments through the glass sphere in the *y*-direction. Each individual beam position simulation was then apertured by the confocal slit (*ω*_*slit*_ = 20 μm), and summed together to yield simulated DSLM and mDSLM images. Each simulation image was normalized by a corresponding simulation in the absence of the glass sphere. For all simulations, an illumination wavelength of *λ*_*ex*_ = 660 nm and fluorescence wavelength of *λ*_*em*_ = 680 nm was used to closely match the experimental conditions.

### Optical phantom experiments

For the first set of experiments, solid agarose phantoms were prepared by dissolving and melting agarose in deionized (DI) water (1% w/v) at 100-deg C on a hotplate with a magnetic stir bar. Once fully dissolved, the agarose was cooled to 60-deg C, and polystyrene beads (*d* = 20 μm, *n*_*sphere*_ = 1.59, Polysciences Inc.) were add and mixed uniformly at a concentration of 0.025% w/v. The agarose solution was then poured into molds, cooled at room temperature, sliced into 1 mm cubes, and index matched overnight to *n* = 1.46 by incubating the phantoms in a mixture of ~60% TDE and 40% DI water. 1 mM Methylene Blue (*λ*_*ex/em*_ = 660/680 nm) was also added to make fluorescent gels. In the second set of experiments, identical agarose phantoms were prepared with smaller polystyrene beads (*d* = 6 μm, *n*_*sphere*_ = 1.59, Polysciences Inc.) at an increased concentration of 0.1% w/v.

For the third set of experiments, agarose phantoms were prepared with fluorescent (*λ*_*ex/em*_ = 660/680 nm, *d* = 7 μm, Invitrogen), rather than non-fluorescent polystyrene beads. Prior to pouring the agarose into a mold (while the mixture was cooled to 40 deg C), 20% v/v Intralipid (Sigma-Aldrich) was added to the mixture to achieve a concentration of 1% v/v Intralipid within the phantom. The size distribution and refractive index of the lipid droplets in Intralipid is well documented, and in general the droplets are sub-micron with a refractive index of *n*_*lipid*_ ~ 1.46 - 1.48, and therefore a reasonable approximation of the Mie and Rayleigh scattering objects in biological tissues [32]. To mimic an optically cleared biological tissue, the agarose phantoms were cleared in the same TDE, DI water, and Methylene Blue mixture to yield a background gel with a refractive index of *n*_*gel*_ = 1.46. The 1 mM background concentration of Methylene Blue also provided a ~1:10 fluorescent to background ratio (relative to the fluorescent beads). For all optical phantom images, a background image was subtracted to account for ambient light contamination. An illumination wavelength of *λ*_*ex*_ = 660 nm was used for all experiments.

### Human breast tissue preparation and imaging

Human breast tissue was obtained through an IRB-approved protocol and the University of Washington Northwest Biotrust (NWBT). Breast tissue was first fixed in 10% formalin for 24 hours. After fixation, the tissue was grossly sliced to a thickness of approximately 1 mm, and passively cleared in a mixture of 60% TDE, 40% DI, and 0.1% v/v Eosin for 24 hours. Imaging was performed using *λ*_*ex*_ = 488 nm to excite the Eosin dye. For all captured images in biological tissues, a background image was subtracted to account for ambient light contamination.

### Mouse small intestine tissue preparation and imaging

Small intestine tissue was obtained from a sacrificed mouse. After removal, the small intestine was fixed in 10% formalin for 24 hours. After fixation, the sample was passively cleared in a mixture of 60% TDE, 40% DI, and 100 μm Acridine Orange. Imaging was performed using *λ*_*ex*_ = 488 nm to excite the Acridine Orange dye. For all captured images in biological tissues, a background image was subtracted to account for ambient light contamination.

### Correction for exponential attenuation of illumination light within samples

For images obtained from the Intralipid-infused agarose gel, as well as both biological specimens, scattering of the illumination light leads to exponential attenuation of the illumination light as a function of *z* in the samples. To better assess the changes in image quality as a function of *z*, images were individually normalized for this exponential attenuation.

The correction procedure consisted of first calculating the median intensity decay as a function of z in the image. This median intensity decay was then smoothed using a 10-pixel window and fit to an exponential decay, 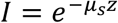, to determine the scattering coefficient of the sample, *μ*_*s*_. For the agarose phantom containing Intralipid, *μ*_*s*_ ~ 10 mm^−1^. In the optically cleared breast tissue, *μ*_*s*_ ~ 1.4 mm^−1^. Finally, in the optically-cleared small mouse intestine, *μ*_*s*_ ~ 0.7 mm^−1^. This is roughly 10 times lower than the scattering coefficient of fresh tissue with no optical clearing (*μ*_*s*_ ~ 10 mm^−1^) [38].

## Data availability

All raw and processed imaging data generated in this work, including the representative images provided in the manuscript and Supplementary Information, are available from the authors upon request.

## Code availability

The custom computer codes used in this study are available from the authors upon request.

## Acknowledgements

Human breast tissues were provided by the NorthWest BioTrust (NWBT), which is supported in part by the NCI (P30CA015704). This work was funded in part by NIH grants F32 CA213615, R01 CA175391, R01 DE023497, the University of Washington Royalty Research Fund, a UW CoMotion Innovation Award, and a Safeway / NCI SPORE developmental award (subcontract) from the Fred Hutchinson Cancer Center. The authors would like to acknowledge and thank Jon Daniels from Applied Scientific Imaging for discussions and consultation regarding the cleared tissue objective.

## Contributions

A.G. and J.L. conceived of the mDSLM concept and designed the studies. A.G., Y.C., C.Y., P.W, and L.B. fabricated the optical setup. A.G. and Y.C. fabricated and imaged the agarose gel samples. A.G. and N.R. prepared and imaged the biological tissues. A.G., Y.C., C.Y., P.W., N.R., and J.L. prepared the manuscript.

